# An explainable machine learning data analytics method using TIGIT-linked genes for identifying biomarker signatures to clinical outcomes

**DOI:** 10.1101/2023.12.05.570321

**Authors:** G Soorya, Divya Agrawal, Shilpa Bhat, Tirtha Mandal, Shalini Kashipathi, M. S. Madhusudhan, Golding Rodrigues, Maloy Ghosh, Narendra Chirmule

**Author notes:** Address of Correspondence: Narendra. Contributed equally to the manuscript.

## Abstract

In the last decade, immunotherapies targeting immune checkpoint inhibitors have been extremely effective in eliminating subsets of some cancers in some patients. Multi-modal immune and non-immune factors that contribute to clinical outcomes have been utilized for predicting response to therapies and developing diagnostics. However, these data analytic methods involve a combination of complex mathematical data analytics, and even-more complex biological mechanistic pathways. In order to develop a method for data analytics of transcriptomics data sets, we have utilized an explainable machine learning (ML) model to investigate the genes involved in the signaling pathway of T-cell-immunoreceptor with immunoglobulin and ITIM domain (TIGIT). TIGIT is a receptor on T, NK, and T-regulatory cells, that has been classified as a checkpoint inhibitor due to its ability to inhibit innate and adaptive immune responses. We extracted gene whole genome sequencing data of 1029 early breast cancer patient tumors, and adjacent normal tissues from the TCGA and UCSC Xena Data Hub public databases. We followed a workflow which involved the following steps: i) data acquisition, processing, and visualization followed by ii) developed of a predictive prognostic model using input (gene expression data) and output (survival time) parameters iii) model interpretation was performed by calculating SHAP (Shapely-Additive-exPlanations); iv) the application of the model involved a Cox-regression model, trained with L-2 regularization and optimization using 5 fold cross validation. The model identified gene signatures associated with TIGIT that predicted survival outcome with a test set with a score of 0.601. In summary, we have utilized this case study of TIGIT-mediated signaling pathways to develop a roadmap for biologists to harness ML methods effectively.

## INTRODUCTION

### Immunotherapy for cancer

In the past decade, there has been a significant decrease in death rates in 11 of the 18 major cancers among men, and 14 of the 20 among women [1-3]. These advances have been attributed to the development and approval of therapies that activate immune pathways [4, 5]. The large effect sizes have been largely due to advances in early detection and more effective treatments. Diagnosis and treatment of cancers has also been transforming from one-size-fits-all (cut-burn-poison) to targeted therapies to more recently algorithm-assisted precision therapies [6]. One of the most significant developments in cancer immunotherapy has been the clinical efficacy of targeting immune checkpoint inhibitors (ICI) such as CTLA-4 and PD-1. However, only 10-30% of patients respond to immunotherapies [7-10]. The limited efficacy has been attributed to several factors, some of which include variability of overall immune health of the patient, varying expression levels of immune checkpoint inhibitors on the tumor and immune cells, differential tumor mutational burden, diversity of immune-modulatory (regulatory, suppressive, and stimulatory) cells and cytokine secretion patterns, genetic and epigenetic factors in both tumor and immune cells, and more recently the microbiome of an individual [11-13]. These results prompt the identification of better biomarkers that can predict immunotherapy response in patients. Therefore, a multi-modal approach to precision immunotherapy is necessary to identify biomarker signatures and improve response to immunotherapy. Such approaches require an analysis of a wide range of data including gene expression analysis, single-cell technologies, imaging modalities [11, 13]. Analysis of these multi-modal data to identify biomarkers requires using ML-based based methods [14].

### Technological advances in data collection and machine-learning based analysis

Advances in technologies have enabled ultra-high throughput data generation, collection, and analysis [15, 16]. **Supplementary Table 1** (available upon request) lists some of the technological advances for collection of data, ML-based algorithms used to analysis the datasets and biomarkers, and multi-modal biomarker signatures identified that correlate with clinical outcomes. Machine learning models such as Support Vector Machines (SVMs), Random Forests, Deep Neural Networks can incorporate several diverse patient characteristics such as clinical attributes, demographics, molecular and immune profiles to generate treatment response models. In addition, tumor characteristics associated with response can also be identified using techniques such as Computer Vision-Aided Radiomics (CVAR) and image analysis to extract tumor features such as size, shape, and heterogeneity. Tumor microenvironments can also be analyzed to determine immune infiltration patterns, spatial distributions and their impact on tumor behavior and treatment response.

In this paper, we chose to work on TIGIT (T-cell immunoreceptor with Ig and ITIM domains) in primary breast cancer datasets, since this receptor is an emerging ICI that has been shown to be significantly upregulated in primary breast cancer tumor tissues [17-19]. TIGIT binding ligands include CD155 and CD112. High TIGIT expression was positively correlated with tumor stage and negatively correlated with progression free survival (PFS) and overall survival (OS) [20]. In an attempt to identify prognostic biomarkers linked with TIGIT, we selected gene sets from public databases. Using protein-protein-interaction (PPI) and network-based computational methods performed network proximity and pathway analysis [14]. We provide a “how-to-guide” of data analytics using machine learning to perform network-biology based identification of gene modules that associated with TIGIT. We have listed activities for data collection, visualization, analysis, interpretation, and application. The conclusion from this analysis is specific to this dataset obtained from the public domain and is intended to be reviewed only for methodology of the data analytics process.

## METHODS

### Case study

*Identifying TIGIT linked genes influencing overall survival in breast cancer patients*. **Figure 1** depicts the steps involved in the data collection, analysis, interpretation.

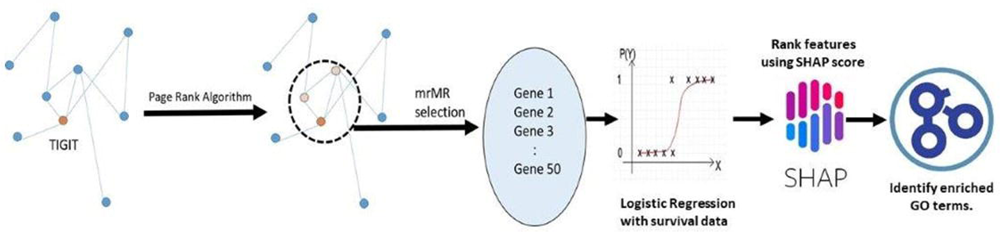

### Data Sets

Genome Data Commons (GDC), The-Cancer-Genome-Atlas (TCGA) and Breast cancer transcriptomics and overall survival datasets were downloaded from UCSC-Xena data hub [21]. From that cohort of patients, the “HTSeq– FPKM” (High-throughput-sequence - Fragments Per Kilo Base Million) data file was downloaded. The expression value for each gene in a given patient was converted from log_2_ (FPKM +1) to FPKM.

### Protein-Protein interaction (PPI) analysis

Using the STRING protein-protein interaction data [22], which was in the form of a list of interacting pairs (protein pairs with interaction score > 700 were only chosen), a graph was constructed for nearly 16k genes using NetworkX package in Python.

### Gene Ranking

The page rank algorithm (link to the code written by netbio on github) was applied to estimate the relative importance of proteins with respect to TIGIT as the central protein of interest. The score quantifying the importance is called *propagation score*. The top 5 percentile of proteins in the list of proteins sorted by propagation score were taken further for analysis. These proteins are members of a sub network relevant to TIGIT function. Nearly 746 were chosen.

### Training the algorithm

Every patient in the survival data set was labeled as “Good surviving” or “Poor surviving” using their overall survival time. The cutoff survival time was chosen in such a manner that the difference between the numbers of dead patients between the resulting two groups is minimal. These labels are to be taken as the ground truth class labels from now on.

The ***train-test split*** was done stratified by the survival label given previously (“Good surviving” vs “Poor surviving”) with 20 percent in test set 80 percent in the training set. Using the train set survival times and dead patient flag, the train data set was split into two sub-groups “Good Prognosis” and “Poor-Prognosis” in a manner similar to the method mentioned previously.

### Feature Selection

For feature selection a version of the “minimum redundance Maximum Relevance” (mrMR) algorithm was implemented as follows: (Figure 2)

1. Using the Good Prognosis and Poor prognosis data subsets derived from the training data, maximum relevance score for each gene feature was computed as the F-statistic computed over the two data subsets.
2. The Redundance score for a gene (Gene_ j) is estimated using the following formula:

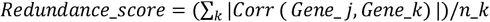

Where the summation is over the set of all genes selected already till the current iteration.
3. A gene is chosen if it maximizes the ratio of Relevance score to Redundance score at each iteration till 50 genes are selected in total.

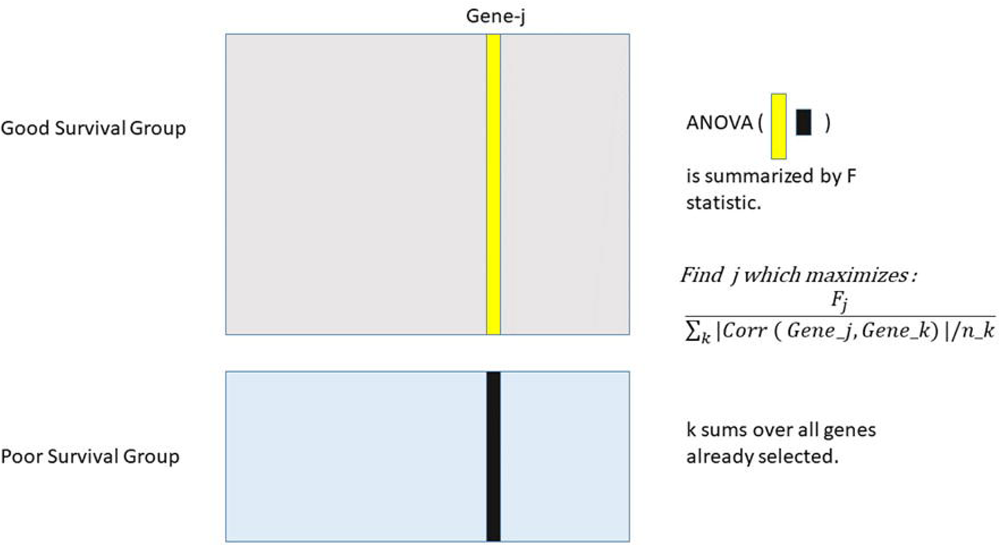

### Cox Regression Analysis

The Cox regression method was analyzed for evaluating the effect of the input transcriptomics data sets, to the overall survival events. This analysis was used to find the statistical correlation of TIGIT-associated gene(s) with higher survival. The statistical correlation of proportional hazard ratio with clinical outcome of overall survival for each of the genes was determined by Wilcoxon test.

## RESULTS

### Visualization of the data set

Figure 3 shows two different methods to visualize the data set. Visualization of large complex data sets in multiple ways is an important first step in the data analysis process. Several resources to enable data visualization are available in the public domain [23, 24].

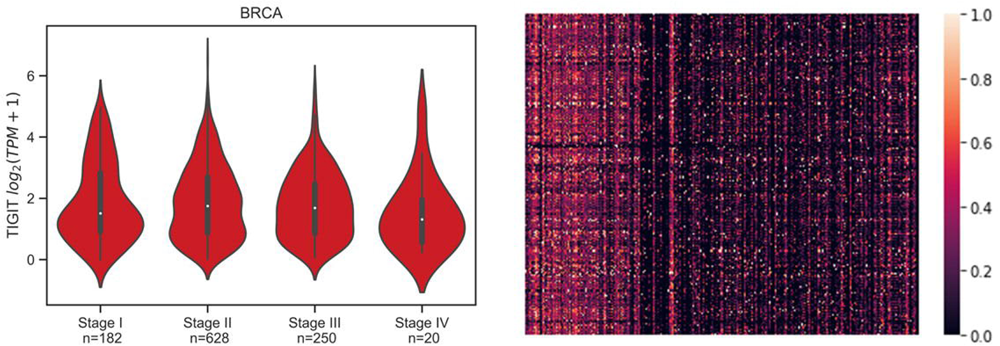

### Testing the mrMR algorithm

The mrMR algorithm was used to identify a 50 gene panel. This was done using the “Good-Prognosis” vs “Poor-Prognosis” sub-groups of the training data (Figure 4a). Then a Logistic regression model with L-2 penalty was trained on the ground truth labels of patients as determined previously using the mrMR selected 50 genes as predictors. 5 fold grid search cross validation was done to optimize the hyper-parameters of each of these models and the best model was saved.

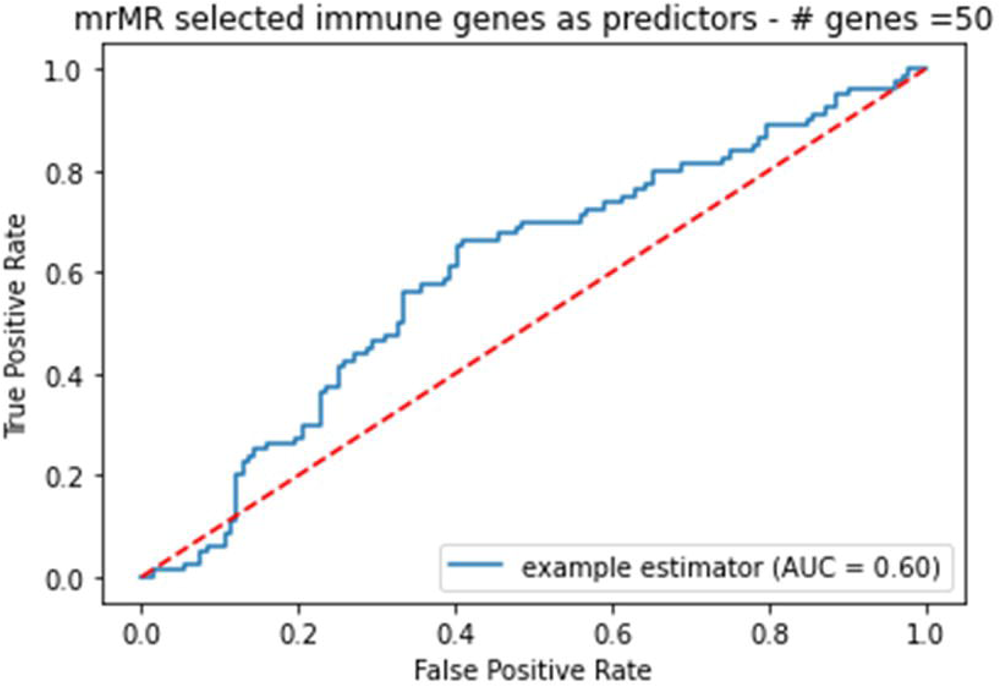

The average absolute SHAP value of each gene feature in the model was computed using the test set (Figure 4b). This score is a measure of the average impact on the model output magnitude. This was then used to rank order genes by their relative importance in contributing to its model’s predictive performance. To estimate the variance of the average absolute SHAP value of each gene feature, the metric was estimated 10 times while keeping the test set constant and summary statistics of the 10 trials were recorded. The top 80 percentile of the genes from the list of 50 genes sorted in descending order by the average absolute SHAP value were used to perform the Gene ontology enrichment analysis for biological process ontology terms using the clusterProfiler package in R. The top ontologies represented in the list were visualized as a dot plot. (Figure 4c) The model identified gene signatures associated with TIGIT that predicted survival outcome with a test set with a score of 0.601 (Figure 4d).

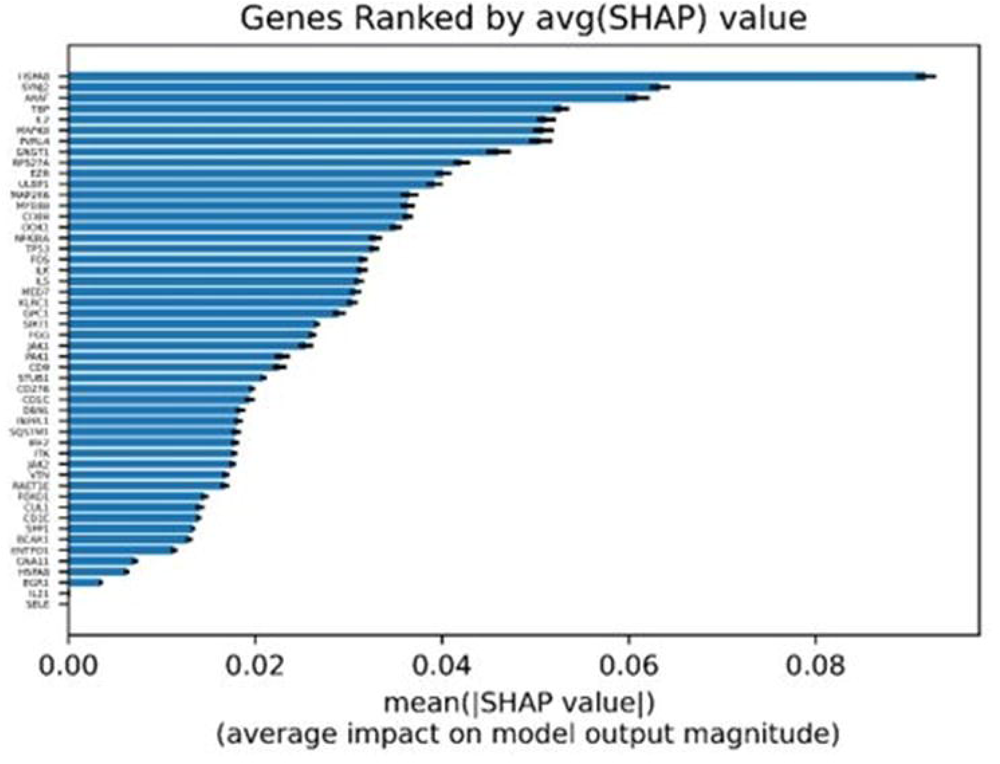

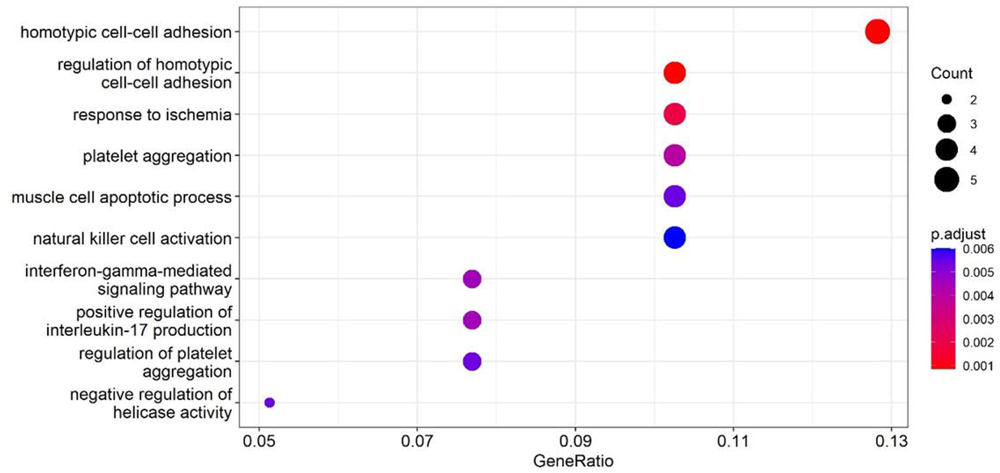

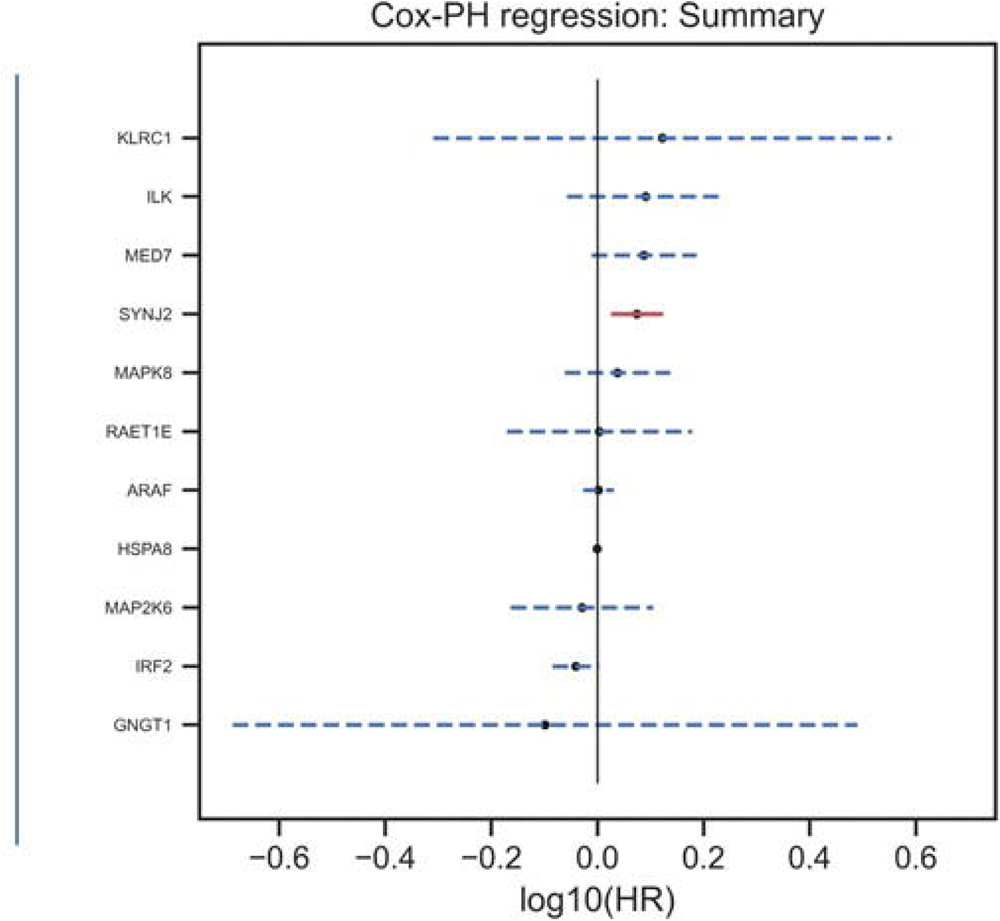

### Cox Regression analysis for identification of the top 10 gene sequences in the signatures associated with TIGIT signaling pathway

The top 10 genes that were associated with better survival in the TIGIT pathway included HSPAB, SYNJ2, ARAF, TBP, IL2, MAPK8, PBRL4, GNGT1, RPS27A, E2R (Figure 5). Based on this data set of input parameters and output outcomes, the model found that the SYNJ2 gene had a significantly increased in the subjects that had a higher survival outcome.

## DISCSSION

In this study, we developed a methodology to evaluate a network-based biomarker(s) that can predict immunotherapy treatment. For conducting this analysis, we used whole genome expression dataset, and survival time data of breast cancer patients from public databases. We focused the analysis on the TIGIT-targeting gene signatures to answer and describe a workflow and methodology to aid biologists collaborating with data scientists.

TIGIT receptors expressed on activated T, NK, T-follicular helper (T-FH) and T-regulatory cells participate in the immunological synapse by interacting with several ligands, e.g., CD112, CD155 (Polio-virus-receptor, PVR, PVRL2)[17, 19, 25]. Several diverse functional responses have been reported upon TIGIT ligation, including cell-adhesion, signal amplification, activation of reactive oxygen species dependent DNA damage responses [26]. TIGIT^-/-^ mice do not develop spontaneous autoimmunity (unlike CTLA4^-/-^ mice) but develop more severe experimental autoimmune encephalitis (EAE), establishing its function as a check-point inhibitor. Signal transduction through TIGIT is mediated through its ITIM and ITT cytoplasmic domain. The cascade of events include modulation of phosphorylation-dephosphorylation equilibrium of proteins such as Grb2, b-arrestin2, phosphatase SHIP-1, inhibiting phosphorylation of ZAP70/Syk, ERK1/3 kinases. These membrane-cytoplasmic events lead to regulation of transcription factors. Engagement of TIGIT – and its ligands has been shown to inhibit immune responses by decreasing IL12 secretion, increased IL-10, reduce the cytolytic capacity of NK cells.

Blockade of TIGIT with PD1 exerts synergistic check-point inhibition that regulates antitumor activities. Several anti-TIGIT monoclonal antibody therapeutics are in clinical development. Etigilimab (OMP-313M32), and tiragolumab (MTIG7192A, RG-6058) are in late stage clinical trials. Recently, tiragolumab in combination with atezolizumab demonstrated an improvement objective response rate and progression-free survival compared with placebo plus atezolizumab in patients with recurrent or metastatic non-small cell lung cancer (NSCLC) [27].

Using the protein-protein interaction STRING database, we identified a list of nearly 16K genes and by assigning a propagation score, defined 746 genes that were used for further analyses. Finally, that analysis of SHAP scores identified the top 80 genes which were assigned based on the closest interactive proteins to TIGIT. The top 10 genes included HSPAB, SYNJ2, ARAF, TBP, IL2, MAPK8, PBRL4, GNGT1, RPS27A, E2R. Kong et al [14] have previously utilized a similar approach to identify PD-1 related gene signatures for bladder, gastric and melanoma tumors. Several of these gene signatures are involved in various aspects of immune regulation such as those for innate immune responses, T cell activation genes, genes involved in signal transduction. Several of these genes have been previously associated with TIGIT mediated signaling pathways.

Mapping the functions of the gene signature associated with TIGIT-mediated signaling pathways will require experimental evidence by obtaining immune and tumor cells from patients treated with anti-TIGIT therapies. Additionally, evaluating this gene signature in patients with breast cancer may provide insights into identifying responder populations. Interestingly, SYNSJ2 (synaptojanin 2), an intracellular protein that participates in membrane trafficking and endocytosis processes, has been associated with turnover of T cell receptor expression [28]. Understanding the roles of these genes may provide clues to development of diagnostics and therapies in for enabling patients’ specific precision medicine.

Taken together, we have described a methodology of data analytics that can be used by biologists to analyze genome sequencing data with clinical outcomes, in order to identify biomarker signature. These signatures could be utilized to develop target-specific diagnostics of disease progression, or companion-diagnostics to monitor effectiveness of immunotherapies and contribute to precision oncology

## Acknowledgement

NC thanks late Jaideep Khare, who had started the work on this manuscript due to his interest and experience in machine learning in drug development.

