## Supplementary Table 1 for "An explainable machine learning data analytics method using TIGIT-linked genes for identifying biomarker signatures to clinical outcomes"

|  | **Application** | **Technologies** |  | **ML Methods** |  | **Databases** |
| --- | --- | --- | --- | --- | --- | --- |
| **Genomics** | DNA sequencing, genetic mutations  Tumor mutational burden (TMB) | Next-generation sequencing |  | Random Forest (RF)  Support Vector Machines (SVM)  Logistic Regression Decision Trees |  | The Cancer Genome Atlas (TCGA)  International Cancer Genome Consortium (ICGC) |
| **Single-cell genomics** |  | ScRNA-Seq | [(https://doi.org/10.1093/pcmedi/pbac002](https://doi.org/10.1093/pcmedi/pbac002) | Drop-Seq, CEL-Seq, Tang method | <https://doi.org/10.1186/s13046-021-01874-1> |  |
| **Chromatin** |  | Chromatin structure Chromatin imaging Long noncoding RNA, DNA-chromatin interaction: 3C, 4C, 5C, ChIP-loop, ChIA-PET, Hi-C, Micro-C XL, CAP-C, Capture Hi-C, HiChIP, GAM, SPRITE, ChIA-Drop, Dip-C DNA-seq, WGS | [https://doi.org/10.1007/s11427-019-1704-2 ,](https://doi.org/10.1007/s11427-019-1704-2) |  |  |  |
| **Transcriptomics** | RNA sequencing | R-Loop-DRIP-seq, R-ChIP bisDRIP-seq, cellular indexing of transcriptomics and epitopes (CITE), microaaray | https://doi.org/10.1016/j.molcel.2019.12.021 (https://doi.org/10.13345/j.cjb.200197 Garcia-Garijo et al., 2019; Zemek et al., 2019) | Differential Gene Expression Analysis Gene Set Enrichment Analysis (GSEA), SVM |  | Gene Expression Omnibus (GEO) TCGA |
| **Epigenetics** | DNA-methylation | Nanopore sequencer, CRISPR-Cas | <https://doi.org/10.1038/s10038-019-0679-0> [https://doi.org/10.1038/s41580-019-0131-5,](https://doi.org/10.1038/s41580-019-0131-5)<https://doi.org/10.1016/j.bbadis.2022.166552> |  |  |  |
| **Single cell- epigenetics** | Tumor infiltrating T cells Single Cell ChIP Seq Single Cell Metabolomics Single Cell Lipidomics Single-cell DNA methylation | Single-cell assay for Transposase-accessible chromatin sequencing (scATAC-seq) Chromatin accessibility: MNase -seq, DNase-seq, FAIRE-seq, Sono-seq, ATAC-seq Flow cytometry, mass spectrometry, Iso-imaging, vibrational spectroscopy, optical imaging | (Blake et al., 2023; Shi et al., 2022) (Ali et al., 2022) |  |  |  |
| **Proteomics** |  | CyTOF | (Helmink et al., 2020) (Gide et al., 2019; Goswami et al., 2020) |  |  |  |
| **Single cell-proteomics** | Single cell mass spectrometry-based proteomics (sco-PE2) | LAESI-MS, MSI | [(https://doi.org/10.1021/jasms.0c00439 )](https://doi.org/10.1021/jasms.0c00439) |  |  |  |
|  | High Dimensional Fluorochrome based flow cytometry (HDF) | SILAC based MS proteomics, TMT-based proteomics | <https://doi-org.ezproxy.library.uq.edu.au/10.1016/j.trac.2020.116005> |  |  |  |
| **Macromolecular Interactomics** | Protein-DNA interaction | ChIP-Seq , ChIP-exo, CUT&Tag, DamID, DamIP, SpDamID, LM-PCR, CE-LIF | <https://doi.org/10.1021/acs.jproteome.1c00074> |  |  |  |
| **Metabolomics** | Single cell Metabolomics | SMALDI, TG-MALDI, NMR based metabolomics, targeted and untargeted metabolomics | <https://doi.org/10.1016/j.drudis.2022.02.018> |  |  |  |
| **Immune Profiling** | Cell types Cell surface receptors Cytokines, Chemokines and solulble factors Signal Transduction second messengers Transcription factors Gene expression | Flow Cytometry Immuno-Histochemistry CyTOF Histopathological analysis T-cell receptor (TCR) sequences B-cell receptor (BCR) sequences ELISA, multiplex immunoassay Smart-Seq 2, multiplex cytokine profiling | ng et al., 2019; Sheih et al., 2020) | RF SVM |  | ImmPort Cancer Immunome Database (TCIA FlowRepository), CRI iAtlas |
| **Radiomics** | Radiographic images | Medical imaging (MRI, CT, PET) |  | CNN Radiometric Analysis |  | TCIA QIN (Quantitative Imaging Network) |
| **Integrated Multi-Omics** | Integrate diverse molecular features to identify predictive biomarkers |  |  | Multi-Kernel Learning, Deep Learning |  | ICGC Human Protein Atlas |
| **Liquid Biopsies** | Circulating tumor DNA (ctDNA) Circulating tumor cells (CTC) | Flow cytometry Cell Sorting Sequencing |  | Deep Learning |  | COSMIC (Catalogue of Somatic Mutations in Cancer CIViC (Clinical Interpretations of Variants in Cancer)) |
| **Clinical Outcome Analyses** |  | Electronic health records Medical Databases |  | Deep Learning (e.g., CNN, RNN) Ensemble Learning methods |  | SEER (Surveillance, Epidemiology, and End Results) NHANES (National Health and Nutrition Examination Survey) |

(References for the TABLE)

li, A., Davidson, S., Fraenkel, E., Gilmore, I., Hankemeier, T., Kirwan, J. A., Lane, A. N., Lanekoff, I., Larion, M., McCall, L.-I., Murphy, M., Sweedler, J. V., & Zhu, C. (2022). Single cell metabolism: current and future trends. *Metabolomics*, *18*(10), 77. <https://doi.org/10.1007/s11306-022-01934-3>

Blake, M. K., O’Connell, P., & Aldhamen, Y. A. (2023). Fundamentals to therapeutics: Epigenetic modulation of CD8+ T Cell exhaustion in the tumor microenvironment. *Frontiers in Cell and Developmental Biology*, *10*. <https://doi.org/10.3389/fcell.2022.1082195>

Da Vià, M. C., Dietrich, O., Truger, M., Arampatzi, P., Duell, J., Heidemeier, A., Zhou, X., Danhof, S., Kraus, S., Chatterjee, M., Meggendorfer, M., Twardziok, S., Goebeler, M.-E., Topp, M. S., Hudecek, M., Prommersberger, S., Hege, K., Kaiser, S., Fuhr, V., … Rasche, L. (2021). Homozygous BCMA gene deletion in response to anti-BCMA CAR T cells in a patient with multiple myeloma. *Nature Medicine*, *27*(4), 616–619. <https://doi.org/10.1038/s41591-021-01245-5>

Deng, Q., Han, G., Puebla-Osorio, N., Ma, M. C. J., Strati, P., Chasen, B., Dai, E., Dang, M., Jain, N., Yang, H., Wang, Y., Zhang, S., Wang, R., Chen, R., Showell, J., Ghosh, S., Patchva, S., Zhang, Q., Sun, R., … Green, M. R. (2020). Characteristics of anti-CD19 CAR T cell infusion products associated with efficacy and toxicity in patients with large B cell lymphomas. *Nature Medicine*, *26*(12), 1878–1887. <https://doi.org/10.1038/s41591-020-1061-7>

Ding, T., & Zhang, H. (2023). Novel biological insights revealed from the investigation of multiscale genome architecture. *Computational and Structural Biotechnology Journal*, *21*, 312–325. <https://doi.org/10.1016/j.csbj.2022.12.009>

Garcia-Garijo, A., Fajardo, C. A., & Gros, A. (2019). Determinants for Neoantigen Identification. *Frontiers in Immunology*, *10*. <https://doi.org/10.3389/fimmu.2019.01392>

Gide, T. N., Quek, C., Menzies, A. M., Tasker, A. T., Shang, P., Holst, J., Madore, J., Lim, S. Y., Velickovic, R., Wongchenko, M., Yan, Y., Lo, S., Carlino, M. S., Guminski, A., Saw, R. P. M., Pang, A., McGuire, H. M., Palendira, U., Thompson, J. F., … Wilmott, J. S. (2019). Distinct Immune Cell Populations Define Response to Anti-PD-1 Monotherapy and Anti-PD-1/Anti-CTLA-4 Combined Therapy. *Cancer Cell*, *35*(2), 238-255.e6. <https://doi.org/10.1016/j.ccell.2019.01.003>

Goswami, S., Walle, T., Cornish, A. E., Basu, S., Anandhan, S., Fernandez, I., Vence, L., Blando, J., Zhao, H., Yadav, S. S., Ott, M., Kong, L. Y., Heimberger, A. B., de Groot, J., Sepesi, B., Overman, M., Kopetz, S., Allison, J. P., Pe’er, D., & Sharma, P. (2020). Immune profiling of human tumors identifies CD73 as a combinatorial target in glioblastoma. *Nature Medicine*, *26*(1), 39–46. <https://doi.org/10.1038/s41591-019-0694-x>

Helmink, B. A., Reddy, S. M., Gao, J., Zhang, S., Basar, R., Thakur, R., Yizhak, K., Sade-Feldman, M., Blando, J., Han, G., Gopalakrishnan, V., Xi, Y., Zhao, H., Amaria, R. N., Tawbi, H. A., Cogdill, A. P., Liu, W., LeBleu, V. S., Kugeratski, F. G., … Wargo, J. A. (2020). B cells and tertiary lymphoid structures promote immunotherapy response. *Nature*, *577*(7791), 549–555. <https://doi.org/10.1038/s41586-019-1922-8>

Ishizuka, J. J., Manguso, R. T., Cheruiyot, C. K., Bi, K., Panda, A., Iracheta-Vellve, A., Miller, B. C., Du, P. P., Yates, K. B., Dubrot, J., Buchumenski, I., Comstock, D. E., Brown, F. D., Ayer, A., Kohnle, I. C., Pope, H. W., Zimmer, M. D., Sen, D. R., Lane-Reticker, S. K., … Haining, W. N. (2019). Loss of ADAR1 in tumours overcomes resistance to immune checkpoint blockade. *Nature*, *565*(7737), 43–48. <https://doi.org/10.1038/s41586-018-0768-9>

Jiang, N., Schonnesen, A. A., & Ma, K.-Y. (2019). Ushering in Integrated T Cell Repertoire Profiling in Cancer. *Trends in Cancer*, *5*(2), 85–94. <https://doi.org/10.1016/j.trecan.2018.11.005>

Rabilloud, T., Potier, D., Pankaew, S., Nozais, M., Loosveld, M., & Payet-Bornet, D. (2021). Single-cell profiling identifies pre-existing CD19-negative subclones in a B-ALL patient with CD19-negative relapse after CAR-T therapy. *Nature Communications*, *12*(1), 865. <https://doi.org/10.1038/s41467-021-21168-6>

Reticker-Flynn, N. E., & Engleman, E. G. (2020). Cancer systems immunology. *ELife*, *9*. <https://doi.org/10.7554/eLife.53839>

Samstein, R. M., Krishna, C., Ma, X., Pei, X., Lee, K.-W., Makarov, V., Kuo, F., Chung, J., Srivastava, R. M., Purohit, T. A., Hoen, D. R., Mandal, R., Setton, J., Wu, W., Shah, R., Qeriqi, B., Chang, Q., Kendall, S., Braunstein, L., … Riaz, N. (2020). Mutations in BRCA1 and BRCA2 differentially affect the tumor microenvironment and response to checkpoint blockade immunotherapy. *Nature Cancer*, *1*(12), 1188–1203. <https://doi.org/10.1038/s43018-020-00139-8>

Sheih, A., Voillet, V., Hanafi, L.-A., DeBerg, H. A., Yajima, M., Hawkins, R., Gersuk, V., Riddell, S. R., Maloney, D. G., Wohlfahrt, M. E., Pande, D., Enstrom, M. R., Kiem, H.-P., Adair, J. E., Gottardo, R., Linsley, P. S., & Turtle, C. J. (2020). Clonal kinetics and single-cell transcriptional profiling of CAR-T cells in patients undergoing CD19 CAR-T immunotherapy. *Nature Communications*, *11*(1), 219. <https://doi.org/10.1038/s41467-019-13880-1>

Shi, P., Nie, Y., Yang, J., Zhang, W., Tang, Z., & Xu, J. (2022). Fundamental and practical approaches for single-cell ATAC-seq analysis. *ABIOTECH*, *3*(3), 212–223. <https://doi.org/10.1007/s42994-022-00082-5>

Xu, Y., Su, G.-H., Ma, D., Xiao, Y., Shao, Z.-M., & Jiang, Y.-Z. (2021). Technological advances in cancer immunity: from immunogenomics to single-cell analysis and artificial intelligence. *Signal Transduction and Targeted Therapy*, *6*(1), 312. <https://doi.org/10.1038/s41392-021-00729-7>

Zemek, R. M., De Jong, E., Chin, W. L., Schuster, I. S., Fear, V. S., Casey, T. H., Forbes, C., Dart, S. J., Leslie, C., Zaitouny, A., Small, M., Boon, L., Forrest, A. R. R., Muiri, D. O., Degli-Esposti, M. A., Millward, M. J., Nowak, A. K., Lassmann, T., Bosco, A., … Lesterhuis, W. J. (2019). Sensitization to immune checkpoint blockade through activation of a STAT1/NK axis in the tumor microenvironment. *Science Translational Medicine*, *11*(501). <https://doi.org/10.1126/scitranslmed.aav7816>
